# A conserved interaction between the effector Sca4 and host endocytic machinery suggests additional roles for Sca4 during rickettsial infection

**DOI:** 10.1101/2024.06.24.600492

**Authors:** Cassandra J Vondrak, Brandon Sit, Chanakan Suwanbongkot, Kevin R. Macaluso, Rebecca L. Lamason

**Affiliations:** Microbiology Graduate Program, Massachusetts Institute of Technology, Cambridge, Massachusetts, USA; Department of Biology, Massachusetts Institute of Technology, Cambridge, Massachusetts, USA; Department of Microbiology and Immunology, Frederick P. Whiddon College of Medicine, University of South Alabama, Mobile, Alabama, USA

## Abstract

Intracellular bacterial pathogens deploy secreted effector proteins that manipulate diverse host machinery and pathways to promote infection. Although many effectors carry out a single specific function or interaction, there are a growing number of secreted pathogen effectors capable of interacting with multiple host factors. However, few effectors secreted by obligate intracellular *Rickettsia* species have been linked to multiple host targets. Here, we investigated the conserved rickettsial secreted effector Sca4, which was previously shown to interact with host vinculin to promote cell-to-cell spread in the model *Rickettsia* species *R. parkeri*. We discovered that Sca4 also binds the host cell endocytic factor clathrin heavy chain (CHC, *CLTC*) via a conserved segment in the Sca4 N-terminus. Ablation of *CLTC* expression or chemical inhibition of endocytosis reduced *R. parkeri* cell-to-cell spread, indicating that clathrin promotes efficient spread between mammalian cells. This activity was independent of Sca4 and appeared restricted to the recipient host cell, suggesting that the Sca4-clathrin interaction also regulates another aspect of the infectious lifecycle. Indeed, *R. parkeri* lacking Sca4 or expressing a Sca4 truncation unable to bind clathrin had markedly reduced burdens in tick cells, hinting at a cell-type specific function for the Sca4-clathrin interaction. Sca4 homologs from diverse *Rickettsia* species also bound clathrin, suggesting that the function of this novel effector-host interaction may be broadly important for rickettsial infection. We conclude that Sca4 has multiple targets during infection and that rickettsiae may manipulate host endocytic machinery to facilitate several stages of their life cycles.

## Introduction

Intracellular bacterial pathogens must adapt to a hostile host cell environment to enable their growth, survival, and dissemination. Consequently, they have evolved various mechanisms to manipulate their host environments to create a more hospitable niche (1–5). One common way for pathogens to interact with host cells is through surface-associated and secreted effector proteins. Bacterial pathogen effectors are known to exert diverse functions, including inducing cellular invasion, establishing vacuolar compartments, remodeling host organelles, modifying the host immune response, and facilitating the spread of these pathogens between cells (6–11).

There is increasing recognition that one efficient way for pathogens to manipulate the host environment is to encode effectors that target multiple proteins or pathways (12–14). For example, the *Salmonella* Typhimurium secreted effector SifA (15, 16) has numerous host targets, binding SKIP (Plekhm2), Plekhm1, and RhoA to support vacuole positioning, integrity, and tubule formation (17–19). The concept of effector multifunctionality is particularly applicable to obligate intracellular bacteria, which depend on their host cells for survival and often have a reduced genome size compared to free-living bacteria or facultative intracellular pathogens (20, 21). One example is the *Anaplasma phagocytophilum* secreted effector Ats-1, which binds to host Beclin-1 to induce the formation of autophagosomes, which are subsequently recruited to the bacterial inclusion to support growth. Ats-1 also localizes to mitochondria, where it is cleaved, and the C-terminal domain blocks apoptosis (22, 23). Therefore, obligate intracellular bacteria are a well-suited model for identifying and characterizing multi-functional effectors, which in turn may provide insight into the full range of host pathways targeted during infection.

*Rickettsia* spp. are a notable group of obligate intracellular pathogens that are transmitted by arthropod vectors (*e.g.*, ticks) to mammals and constitute an emerging public health threat (24, 25). Members of the spotted fever group *Rickettsia*, including the model species *Rickettsia parkeri*, are a pathogenic subset of the genus that undergo a unique life cycle. They must invade a host cell, escape from the vacuole, and replicate in the cytosol. Subsequently, they hijack host actin to become motile and eventually, they manipulate the host membrane to move from one cell to the next in a process called cell-to-cell spread (26, 27).

Only a few secreted effectors have been functionally characterized across the genus (26, 28, 29). One such partially characterized effector is surface cell antigen 4 (Sca4), a large 120kDa protein that despite its name, is secreted into host cells, where it interacts with the host adhesion protein vinculin through two distinct vinculin-binding sites (30, 31). Prior work using a *R. parkeri* transposon mutant lacking Sca4 (*sca4*::Tn) showed that Sca4 promotes cell-to-cell spread by competitively disrupting the normal vinculin-α-catenin interaction at adherens junctions to tune intercellular tension (32). Sca4 is well conserved across the *Rickettsia* genus, even in species not thought to undergo cell-to-cell spread (28, 31, 33–35). However, beyond its interaction with vinculin, Sca4 has no other known targets during infection in either mammals or in arthropod vectors.

To better understand the role of Sca4 during infection, we investigated whether Sca4 has any additional host-binding partners and if they too promote Sca4’s role in cell-to-cell spread. We discovered that Sca4 interacts with the canonical endocytosis protein clathrin heavy chain and that the amino terminus of Sca4 is necessary and sufficient for this interaction. Intriguingly, we found that clathrin supports cell-to-cell spread independently of Sca4, suggesting that Sca4 and clathrin have multiple roles in infection.

## Results

### Sca4 interacts with clathrin heavy chain

Beyond its well-defined vinculin-binding sites, the remainder of the *R. parkeri* Sca4 sequence has not been functionally characterized, despite its large size (1023 aa). We hypothesized that Sca4 could be a candidate multi-functional effector with additional host interacting partners. We immunoprecipitated Sca4 from lysates of A549 cells infected with wild-type (WT) *R. parkeri* and first looked for the presence of obvious interacting partners via SDS-PAGE and Coomassie staining. Here, two prominent bands were apparent, which we excised and identified via mass spectrometry. As expected, the 120 kDa band corresponded to Sca4, while the ~190 kDa band was identified as the host protein clathrin heavy chain (CHC, *CLTC*) (Fig. 1A). Clathrin heavy chain is an essential protein that supports clathrin-mediated endocytosis (CME) from the plasma membrane and trafficking throughout the cell, including secretion from the trans-Golgi network and recycling from early endosomes (36). Because no other bands were clearly visible with this approach, we repeated this IP in triplicate and analyzed the samples via LC-MS/MS to more broadly capture the Sca4 interactome. The dominant interaction partner for Sca4 across all replicates was CHC (Fig. 1A). However, we did not identify any additional endocytic proteins that interact with Sca4, including clathrin light chain (Fig. 1B). We also did not observe vinculin, which was expected as crosslinking conditions are usually required to co-immunoprecipitate vinculin (37–39). We also noted that the human plasma protein histidine-rich glycoprotein (HRG) was enriched in the infected sample. As HRG interacts with IgG and is upregulated during infection (40), we suspected this was an artifact and focused our efforts on studying the novel Sca4-clathrin interaction.

**Figure 1.**
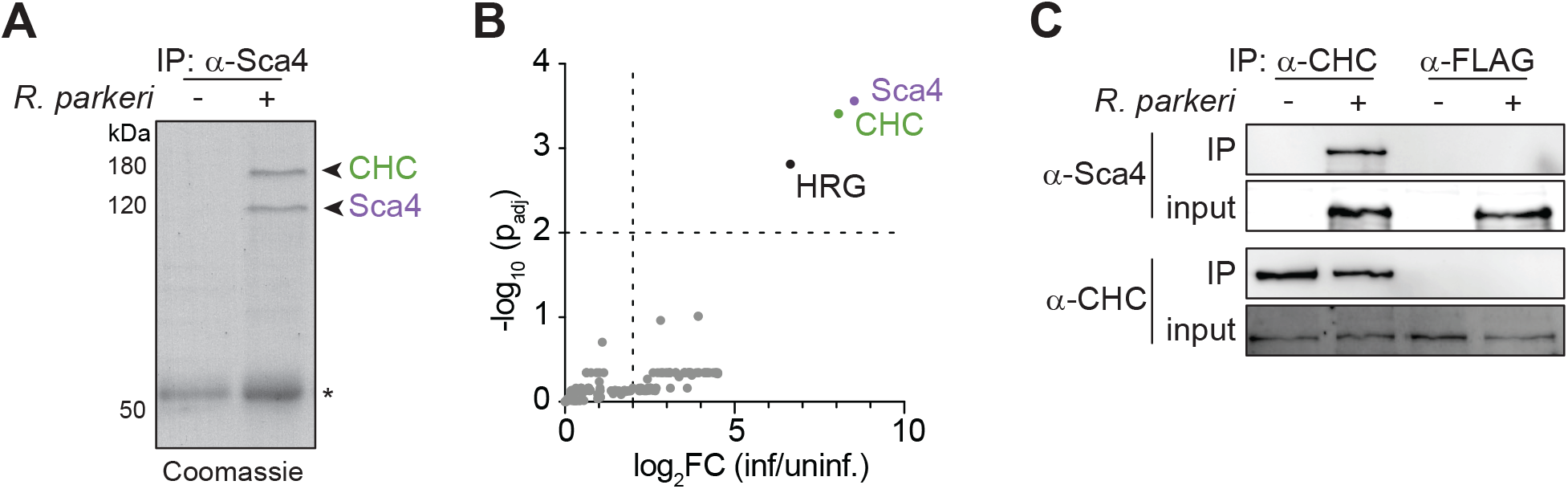
Sca4 interacts with clathrin heavy chain. (A) Sca4 IP from uninfected or WT *R. parkeri*-infected A549 cell lysates, with samples visualized on a Coomassie-stained SDS-PAGE gel. Band identities for Sca4 and CHC were confirmed by mass spectrometry. Asterisk indicates antibody heavy chain. (B) Enriched proteins from IP-MS of Sca4 from uninfected or WT *R. parkeri*-infected A549 cell lysates. Fold change of protein abundance and p_adj_ (unpaired two-tailed t-test, df = 4, with 1% false discovery rate adjustment) were calculated for n = 3 independent samples. Sca4, CHC (clathrin heavy chain), HRG (histidine-rich glycoprotein) and thresholds of fold change > 4 and p_adj_ < 0.01 are indicated. (C) Western blot following CHC or FLAG IP from WT *R. parkeri*-infected or uninfected A549 cell lysates.

To validate the Sca4-clathrin interaction, we performed the reciprocal IP by immunoprecipitating endogenous clathrin from *R. parkeri*-infected host cell lysates and probing for Sca4. We observed the interaction between CHC and Sca4 with this approach and not when a control FLAG-specific antibody was used (Fig. 1C). Together, these results demonstrate that Sca4 and clathrin interact during *R. parkeri* infection.

### The N-terminal domain of Sca4 interacts with clathrin

Given the robust interaction observed between Sca4 and clathrin, we wondered if Sca4 interacted with CHC via a canonical clathrin-binding motif, but bioinformatic and manual screening of known motifs did not yield any promising candidate regions (41, 42). To experimentally define the region of Sca4 that is responsible for the interaction with CHC, we first built a series of 3xFLAG-tagged Sca4 domain truncations (Fig. 2A domain diagram). Domains were predicted by comparing the Sca4 secondary structure predictions from DomPred (43) with the published partial crystal structure (44). We then transiently expressed these truncations in HEK293T cells and used a co-IP assay to measure their interactions with endogenous clathrin. We observed that Sca4 truncations lacking the first N-terminal domain of Sca4 (Sca4-D1) did not interact with endogenous clathrin, suggesting that Sca4 domain 1 (Sca4-D1) is necessary for the clathrin interaction (Fig. 2B). Importantly, the absence of domain 1 did not abolish the interaction of Sca4 with vinculin (Fig. S1). We also noted that clathrin levels were lower in the samples expressing full-length Sca4 (Fig. 2B), which aligns with prior work showing that the overexpression of Sca4 can be toxic (32).

**Figure 2.**
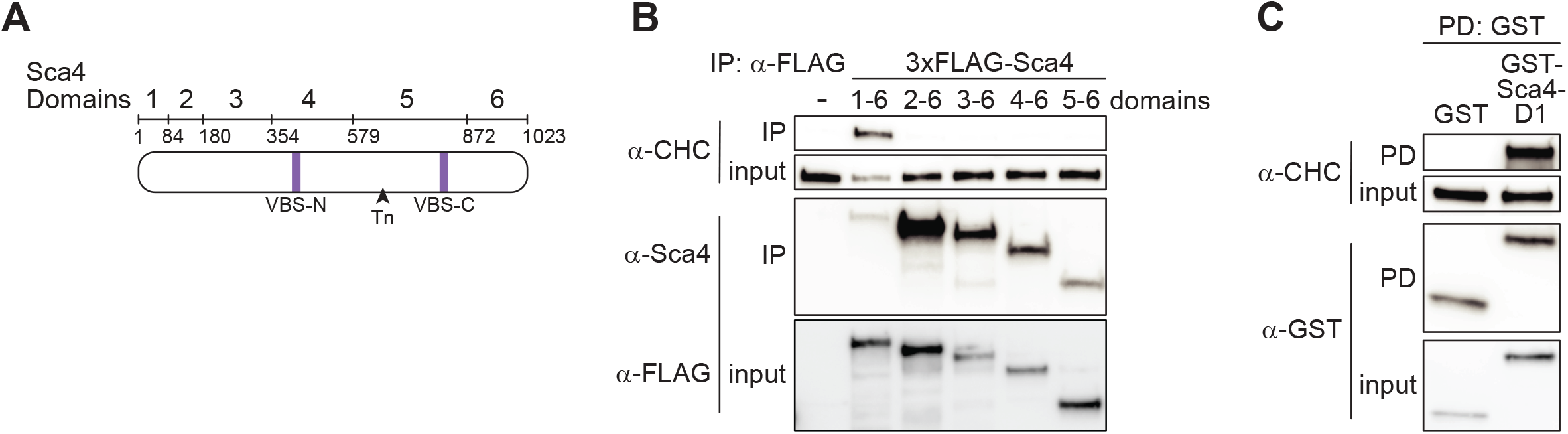
The Sca4 N-terminus interacts with clathrin heavy chain. (A) Predicted Sca4 domains in *R. parkeri* used for Sca4 truncation experiments. Vinculin binding sites (purple) and transposon insertion site for *sca4*::Tn (triangle) are marked. (B) Western blot of the co-IP between the 3x-FLAG-Sca4 truncations transiently expressed in HEK293T cells and endogenous CHC. (C) Western blot for CHC and glutathione-S-transferase (GST) following GST pull-down (PD) of transiently expressed GST-Sca4-D1 or GST-only control in HEK293T lysates.

To test whether domain 1 was sufficient for the Sca4-clathrin interaction, we transiently expressed GST-tagged Sca4-D1 or GST alone in HEK293Ts and performed pulldown assays. We detected a robust interaction between GST-Sca4-D1 and endogenous clathrin but not the GST-only control (Fig. 2C). Together, these results show that domain 1 of Sca4 is both necessary and sufficient to interact with CHC.

### Clathrin contributes to efficient *R. parkeri* cell-to-cell spread

Our prior work showed that the primary infection defect of *sca4*::Tn *R. parkeri* in mammalian cells is in cell-to-cell spread, and not earlier stages of infection such as invasion, actin-based motility, or bacterial replication (32). This led us to ask whether clathrin may also promote spread. We tested this hypothesis by silencing *CLTC* expression via RNAi, and measuring cell-to-cell spread efficiency using an infectious focus assay. In this assay, host cell monolayers are infected with *R. parkeri* at a low multiplicity of infection (MOI) to create spatially isolated infection foci established from a single invasion event. Spread efficiency is then quantified by counting the number of infected cells in each focus at 28 hours post-infection (hpi). siRNA-mediated depletion of CHC reduced WT *R. parkeri* spread (Fig. 3A, C), suggesting that clathrin promotes the spread of *R. parkeri*. This reduction was similar in magnitude to the spreading defect seen with the *sca4*::Tn mutant strain (32), and a reduction in CHC expression did not further enhance the mutant phenotype (Fig. 3A). We also counted the number of bacteria per infectious focus and noted that one of the siRNA treatments correlated with a small but statistically significant reduction in bacterial counts (Fig. 3B). Because clathrin also promotes rickettsial invasion (45), it is possible that infecting cells with reduced CHC levels will delay invasion and slightly alter infection dynamics.

**Figure 3.**
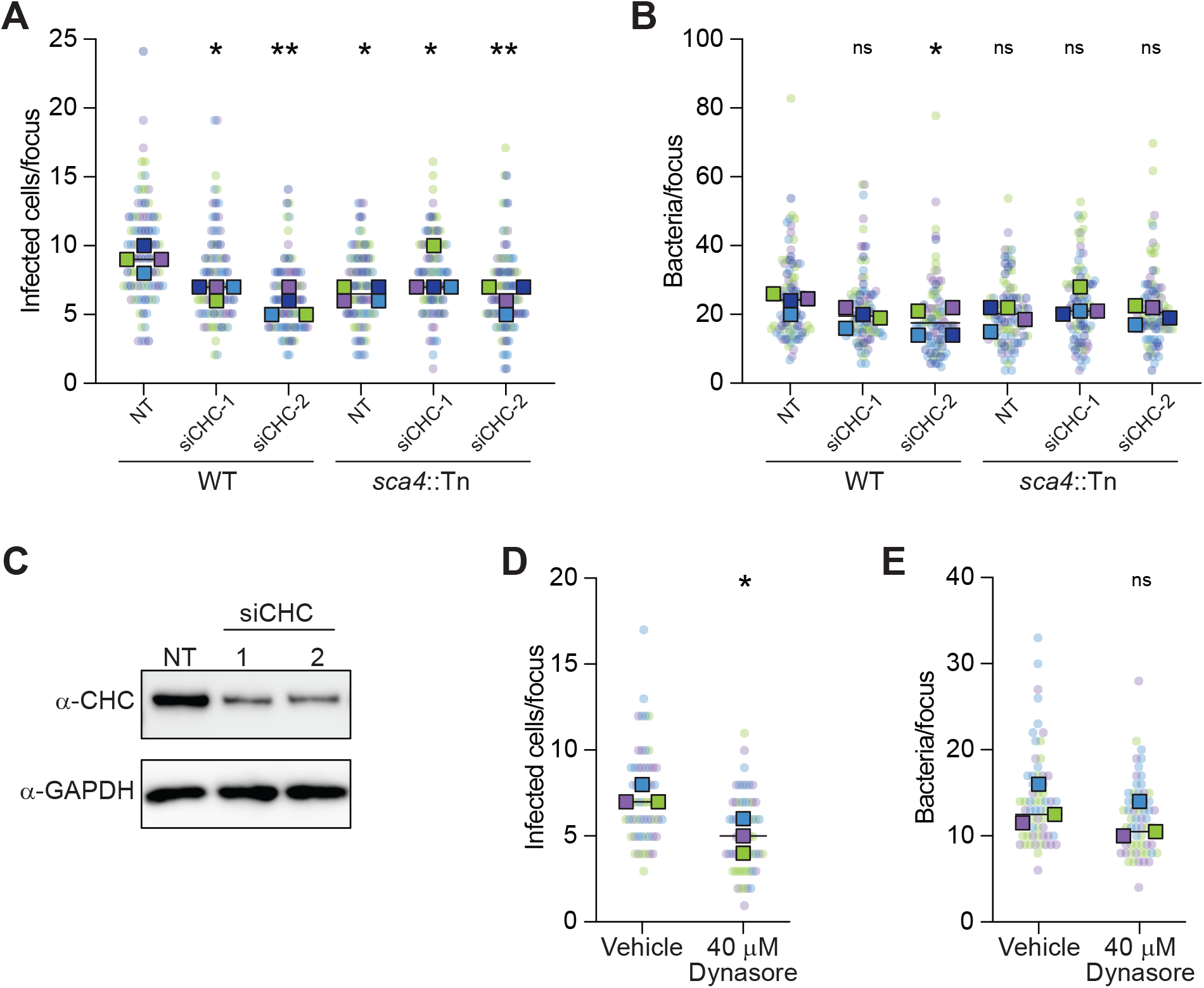
Clathrin promotes *R. parkeri* cell-to-cell spread. (A) Infectious focus assay results for WT or *sca4*::Tn *R. parkeri* at 28 hpi after siRNA-mediated silencing of *CLTC* (siCHC-1, siCHC-2) vs a non-target control (NT). Medians from n = 4 independent experiments (squares) are superimposed over the raw data (circles) and were used to calculate medians (black line) and p-values relative to NT-treated WT infection (Ordinary one-way ANOVA with uncorrected Fisher’s LSD post-hoc test, *p < 0.05, **p < 0.005, ***p < 0.0005). Data are shaded by replicate experiment. (B) Bacteria counts showing total bacteria per focus from the infectious focus assay as described in (**A**). ns = not statistically significant (p > 0.05). (C) Western blot of CHC and GAPDH (loading control) from A549 lysates with indicated siRNA treatment from (**A**, **B**). (D) Infected cells per focus from an infectious focus assay treated with 40 μM dynasore or vehicle (EtOH) during the last 8 hours of infection with WT *R. parkeri*. Medians from n = 3 independent experiments (squares) are superimposed over the raw data (circles) and were used to calculate medians (black line) and p-values (unpaired two-tailed t-test, *p < 0.05). Data are shaded by replicate experiment. (E) Bacteria counts showing total bacteria per focus from the infectious focus assay as described in (**D**).

As an orthogonal approach, we tested clathrin’s role in spread by treating pre-infected (i.e., post-invasion) cells with the small molecule dynasore to inhibit endocytosis. Dynasore targets dynamin, which is critical for the scission of endocytic vesicles during CME (46). To avoid interfering with *R. parkeri* host cell invasion and to mitigate any long-term toxic effects of dynasore treatment, we measured spread in monolayers treated with 40µM of dynasore for only the last 8 hours of infection. We confirmed that this treatment regime inhibited clathrin-mediated endocytosis using a transferrin uptake assay. Dynasore-treated cells took up significantly less fluorescently labeled transferrin than a vehicle-only control, indicating successful inhibition of CME (Fig. S2). Under these same conditions, we observed a marked reduction in WT *R. parkeri* spread, but not growth, after dynasore treatment (Fig. 3D, 3E). Together, these results show that clathrin promotes efficient WT *R. parkeri* spread.

### The Sca4-clathrin interaction is dispensable for spread

Our prior work showed that Sca4 specifically acts in the donor host cell to promote spread into the neighboring recipient cell (32). If Sca4 interacts with clathrin to promote spread, we predicted this would also have to occur in the donor cell. To test this hypothesis, we first transformed the *sca4*::Tn *R. parkeri* mutant with a plasmid stably expressing FLAG-tagged full-length Sca4 (32) or ΔD1-Sca4. To confirm that FLAG-ΔD1-Sca4 was secreted from *R. parkeri*, we used selective lysis after infection to separate the host cytoplasm, which contained secreted effectors like Sca4, from the intact bacteria. FLAG-ΔD1-Sca4 and FLAG-Sca4 were readily detected in the secreted and pellet fractions, indicating that the deletion of D1 does not impair secretion from or expression in *R. parkeri*, respectively (Fig. 4A). We then used a mixed-cell spread assay to determine if the Sca4-clathrin interaction was necessary for spread from the donor cell. In this assay, a small number of fluorescently labeled pre-infected “donor” cells are mixed with a population of unlabeled “recipient” cells. Spread is measured by calculating the percent of bacteria that spread from the donor cell to the surrounding recipient cells. Consistent with our prior observations, the expression of full-length Sca4 complemented the spread of the *sca4*::Tn mutant (32). Surprisingly, we found that ΔD1-Sca4 also rescued the *sca4*::Tn mutant spread defect to the same level as the full-length Sca4 (Fig. 4B). These data suggest that while clathrin promotes WT spread, the Sca4-clathrin interaction is dispensable for efficient cell-to-cell spread.

**Figure 4.**
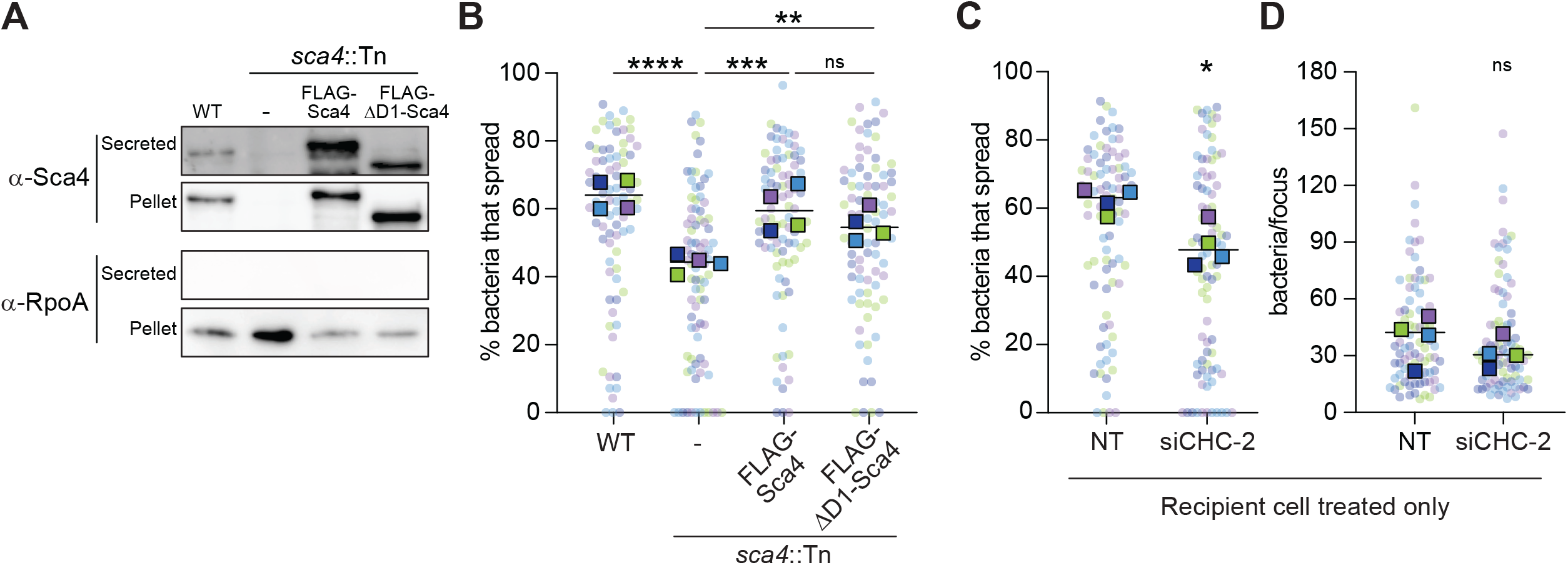
Clathrin acts independently of Sca4 in recipient cells to promote spread. (A) Western blot of Sca4 and RpoA from selective lysis of A549 cells infected with the indicated *R. parkeri* strains. Secreted samples represent protein in host cytoplasm, while pellets contain intact bacteria. RpoA (RNA polymerase subunit α), bacterial lysis control. (B) Mixed-cell spread assay showing the percent of bacteria that spread from a donor cell comparing infections with strains tested in (**A**). Medians from n = 4 independent experiments (squares) are superimposed over the raw data (circles) and were used to calculate medians (black line) and p-values (Ordinary one-way ANOVA with uncorrected Fisher’s LSD post-hoc test *p < 0.05, **p < 0.005, ***p < 0.0005, ****p < 0.0001). Data are shaded by replicate experiment. (C) Mixed-cell spread assay with WT *R. parkeri* after silencing *CHC* expression in the recipient cells only. *p < 0.05 (unpaired two-tailed t-test of biological replicate medians) (D) Total bacteria per infectious focus for the mixed cell assay described in (**C**). ns = not statistically significant (p > 0.05).

We wondered if clathrin supported efficient spread specifically in the recipient rather than donor cell, which could explain why the Sca4-clathrin interaction did not influence spread. Similar results have been observed for *Shigella flexneri*, which relies on CME from the recipient cell to promote cell-to-cell spread (47). To test this hypothesis, we used the mixed-cell spread assay to measure spread into recipient cells treated with either a non-target control or *CLTC*-specific siRNA. This experimental setup avoids altering clathrin-mediated invasion into donor cells by only reducing clathrin expression in the recipient cells. We detected a significant reduction in WT spread when *CLTC* expression was reduced in the recipient cells (Fig. 4C) without significant changes in rickettsial burdens (Fig. 4D). These data are consistent with the idea that clathrin promotes spread by acting in the recipient cell through a Sca4-independent mechanism.

### Sca4 does not alter host transferrin trafficking

Given the spread-independent role of the Sca4-clathrin interaction, we hypothesized that the Sca4-clathrin interaction might impact key cell-intrinsic, clathrin-dependent functions like CME and recycling. We first tested whether ectopic expression of Sca4 was sufficient to alter CME kinetics. We used A549 cells stably expressing Sca4 or an empty vector control (32) and measured endocytosis via transferrin uptake assays. Transferrin is an iron-scavenging glycoprotein that binds the transferrin receptor, which in turn is endocytosed exclusively by CME, making it the canonical receptor for CME kinetic studies (48, 49). There was no discernable difference in transferrin uptake between the Sca4-expressing cells and controls, suggesting that ectopic expression of Sca4 was not sufficient to alter CME of transferrin (Fig S3A). To complement this approach, we also performed the transferrin uptake assay on cells infected with WT or *sca4*::Tn *R. parkeri.* However, no major or consistent differences in transferrin uptake were observed across all timepoints (Fig. S3B), indicating that infection does not grossly impact CME of transferrin.

Clathrin is also involved in transferrin recycling from recycling endosomes to the plasma membrane (50, 51). We used the A549 cells from above that stably expressed Sca4 or an empty vector control to test if Sca4 impacted transferrin recycling rates. As above, no difference was observed for clathrin-mediated recycling of transferrin between Sca4 and control cells (Fig. S3C). These data indicate that Sca4 does not alter the CME or recycling of transferrin in host cells, although it does not eliminate the possibility that other ligands or receptors are affected.

### The Sca4-clathrin interaction is conserved within the *Rickettsia* genus and required for optimal growth in tick cells

Our data suggest that Sca4 supports another aspect of infection beyond cell-to-cell spread. Sca4 is present across the *Rickettsia* genus (33), even in members not thought to undergo cell-to-cell spread like *R. prowazekii* and *R. typhi*, (31, 35). However, no other roles have been described for Sca4. To determine if the Sca4-clathrin interaction was conserved across the genus, we first examined the sequence conservation of the D1 region across a select number of *Rickettsia* species. These representatives, though not exhaustive, highlight members in the spotted fever group (*R. parkeri*, *R. rickettsii*, and *R. conorii*), the typhus group (*R. prowazekii* and *R. typhi)*, and the non-pathogenic species *R. bellii*. The D1 regions of all species tested were predicted by AlphaFold and DomPred to have disordered regions (Fig. 5A, dotted line) flanking a short alpha-helix (Fig. 5A). Within the D1, high sequence similarity was noted within this helix and the sequence immediately upstream, suggesting that the interaction may be conserved. To test this hypothesis, we transiently expressed GST-Sca4-D1 fusions from three diverse *Rickettsia* species in HEK293T cells and performed pulldown assays. In agreement with our prediction, all three D1 sequences interacted with endogenous CHC (Fig. 5B).

**Figure 5.**
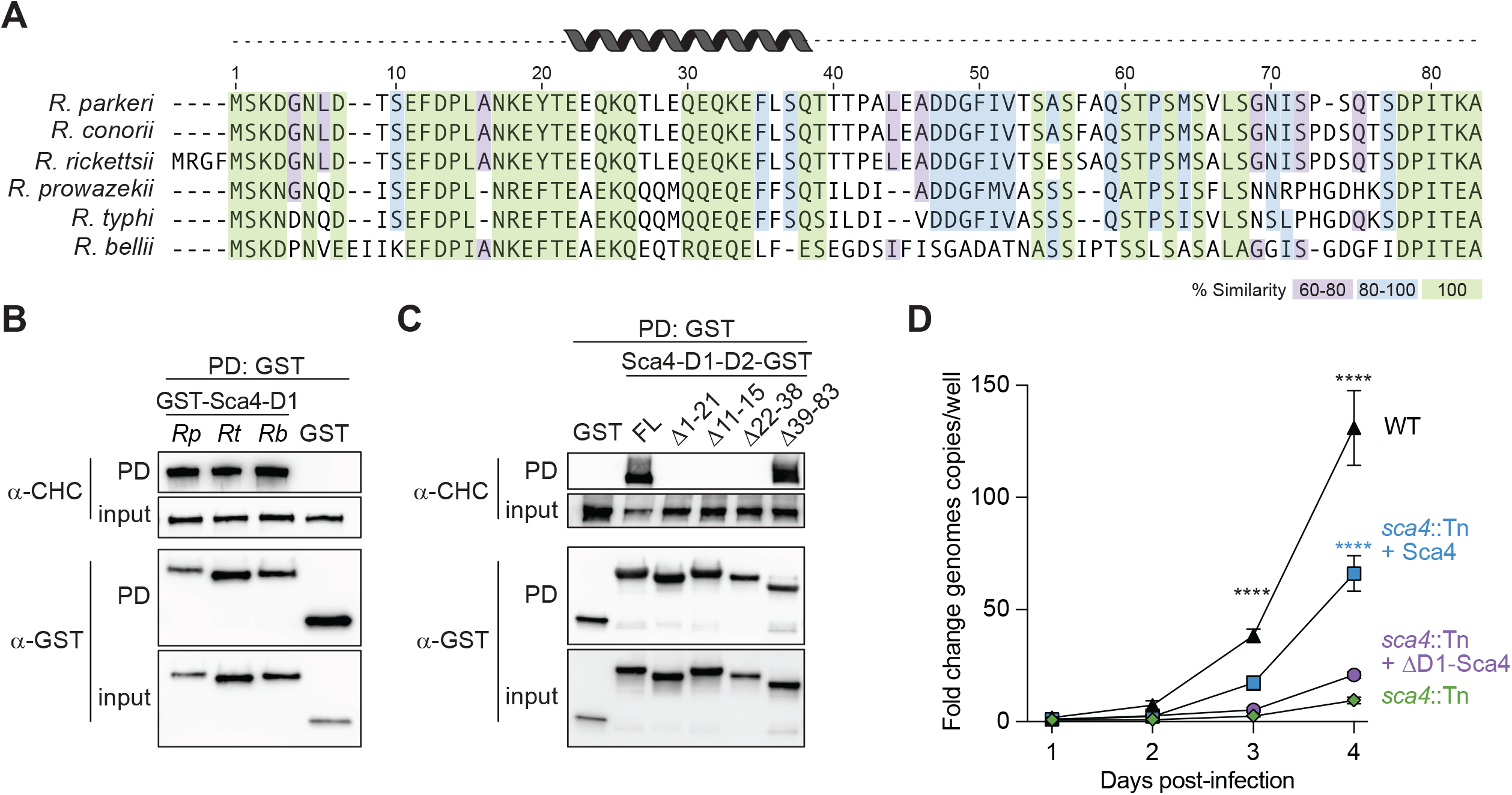
The Sca4-clathrin interaction is conserved among *Rickettsia* species and required for *R. parkeri* growth in tick cells. (A) Alignment of Sca4-D1 sequences across selected *Rickettsia spp*. Colors denote percent amino acid similarity between species. Predicted secondary structure is shown. (B) Western blot for CHC and GST following GST pull-down of transiently expressed GST-Sca4-D1 from *R. parkeri* (*Rp*), *R. typhi* (*Rt*), and *R. bellii* (*Rb*) and a GST only control in HEK293T cells. (C) Western blot for CHC and GST following a GST pull-down of transiently expressed Sca4-D1-D2-GST truncations in HEK293T lysates. FL, full-length D1-D2 construct. Numbers denote the amino acids deleted (Δ) in each truncation. (D) Growth kinetics of indicated *R. parkeri* strains in ISE6 tick cells with mean ± SEM from two independent replicates, each with three technical replicates, normalized to bacteria input 1 hpi. Two-way ANOVA with Tukey’s multiple comparisons test was performed to determine the statistical significance compared to *sca4*::Tn loads at a given time point (*p < 0.05, ****p < 0.0001).

Given this conservation, we hypothesized that the regions of highest similarity would be necessary for the Sca4 interaction and tested this idea by making truncations of the *R. parkeri* D1 region. To facilitate their expression in HEK293T cells, we transiently expressed these truncation mutants as C-terminally tagged D1-D2-GST fusions and performed pulldown assays. While the D1-D2-GST fusion readily pulled down endogenous clathrin, deletion of the residues corresponding to the conserved helix and upstream disordered region completely abolished the interaction. Deletion of the residues downstream of the helix did not impair the interaction (Fig. 5C). Thus, the Sca4-clathrin interaction is maintained in diverse members of the *Rickettsia* genus and is dependent on a highly conserved region of the D1 domain.

All members of the *Rickettsia* genus inhabit arthropod vectors, where they maintain a population through transstadial and transovarial transmission (52, 53). Given the conservation of Sca4, we wondered if Sca4 contributed specifically to infection of the arthropod vector. We found that unlike in mammalian cells, *sca4*::Tn mutant has reduced bacterial burdens in ISE6 tick cells. This phenotype is partially rescued by a full-length Sca4 but not the ΔD1-Sca4 construct, suggesting that Sca4-D1 is necessary for productive infection in tick cells (Fig. 5D). Moreover, these data suggest that Sca4 regulates different aspects of the infectious life cycle in a cell type-specific manner.

## Discussion

With a limited arsenal of effectors at their disposal, some intracellular pathogens encode proteins capable of acting on multiple host targets. For obligate intracellular bacteria with severely reduced genomes, this strategy is likely quite advantageous, but so far poorly understood. Here, we discovered that the *R. parkeri* effector Sca4 has multiple targets as we found a novel interaction between Sca4 and clathrin heavy chain. We demonstrate that clathrin, a key host endocytosis and trafficking protein, is required for efficient cell-to-cell spread but does so independently of its interaction with Sca4. We propose that clathrin plays multiple, spatially segregated roles in infection - one in the recipient cell to promote spread, and another possibly conserved Sca4-dependent role in the donor cell. Our work further suggests that the Sca4-clathrin interaction supports bacterial growth in tick cells, highlighting the distinct targets and functions of Sca4 across different host and vector environments.

Clathrin and CME are common targets for pathogens (54, 55). Many viruses (VSV, Ebola (56, 57)), bacteria (e.g., *R. parkeri* and *Listeria monocytogenes* (45, 58)), and eukaryotic pathogens (e.g. *Candida albicans* and *Trypanosoma cruzi* (59, 60)) depend on clathrin to invade host cells. Once inside the cell, some bacterial pathogens have evolved additional strategies to hijack clathrin function. For example, *Coxiella burnetii* recruits clathrin to the *Coxiella*-containing vacuole (CCV) to facilitate proper CCV expansion and intracellular replication (61). Clathrin also supports the dissemination of pathogens like *S. flexneri*. Similar to our study with *R. parkeri*, *S. flexneri* relies on clathrin in the recipient host cell to promote cell-to-cell spread (47). Clathrin is the first trafficking protein to be implicated in *R. parkeri* spread and further work will be needed to fully understand how it supports spread.

Bacteria often use effectors to co-opt clathrin pathways indirectly by targeting clathrin adaptors and accessory proteins. In *C. burnetii*, the effector CvpA contains sorting motifs that interact with the adaptor protein AP2 to promote clathrin enrichment at the *Coxiella-*containing vacuole (CCV) and promote bacterial growth (61). Another *C. burnetii* effector Cig57 uses an endocytic sorting motif to interact with FCHO2 to facilitate the formation of the CCV (62). Recent work showed that the *Salmonella* Typhimurium effector SpvB interacts via IP with clathrin and the adaptor AP1 to slow endocytosis and disrupt protein secretion (63). Unlike these other effectors, Sca4 does not contain an obvious clathrin binding motif, and we were unable to detect other clathrin adaptor proteins in our IP-MS data. Thus, we propose that Sca4 directly interacts with clathrin, likely through a novel clathrin binding motif. However, as we did not detect clathrin light chain in our IP-MS results, our list of interactors may be incomplete, and we cannot eliminate the possibility that the Sca4-CHC interaction is facilitated by another protein. If the interaction between Sca4 and clathrin is direct, then it likely relies on a novel candidate clathrin binding motif within the highly conserved N-terminal residues of Sca4. Continued investigation will be needed to test this hypothesis and to map which region of clathrin is also required for the Sca4-CHC interaction.

We did not detect a Sca4-dependent impact on CME and recycling of transferrin in human cells, suggesting Sca4 does not globally alter clathrin-mediated trafficking. However, we cannot eliminate the possibility that Sca4 alters clathrin-mediated trafficking of only a subset of host proteins or that CME-dependent pathways are specifically targeted in tick cells. Alternatively, the Sca4-clathrin interaction could alter a non-endocytic cellular process regulated by clathrin to impact intracellular bacterial infection. For instance, clathrin stabilizes the mitotic spindle to ensure proper chromosome segregation (64, 65) and clathrin-mediated trafficking from the Golgi influences many cell functions from the maintenance of polarity (66) to the presentation of MHC-II (67). Clathrin can even participate in immune regulation by controlling the baseline activation of NF-κB (68). As *R. parkeri* remodels the cell into an ideal intracellular niche, tuning one or more of these processes could be advantageous.

Considering the known role of Sca4 in promoting cell-to-cell spread, we were initially surprised to find that the Sca4-clathrin interaction does not promote cell-to-cell spread in mammalian cells. However, considering Sca4’s conservation throughout the *Rickettsia* genus (33) and evidence that Sca4 is under positive selection (34), even in species not thought to undergo cell-to-cell spread, we speculate that the Sca4-clathrin interaction may promote infection through a pathway or life cycle stage that is also conserved across the genus. This idea is supported by our finding that Sca4 is critical for *R. parkeri* growth in tick cells, since all *Rickettsia* spp. interface with ticks or other arthropods in the environment. As the loss of Sca4 is not associated with a similar growth defect in mammalian cells (32), further work is needed to fully understand the molecular basis of host and vector specificity in Sca4’s contribution to rickettsial fitness.

Recent work has highlighted the value of understanding effector function in both vector and host as effectors may support different functions in these species. For example, the rickettsial surface protein rOmpB modulates the growth of *R. parkeri* in mammalian, but not tick cells (69). In contrast, the rickettsial effectors Sca2 and RickA both support actin-based motility and cell-to-cell spread in tick and mammalian cell lines (70). Thus, it will be important to further examine the role of Sca4 in arthropod cells and determine to what extent this role is dependent on clathrin, which is conserved across eukaryotes (71) and essential in ticks (72).

Investigating the full interactome of known secreted effectors is critical to better understanding *Rickettsia* spp. pathogenesis. Our work illustrates how a key rickettsial secreted effector, Sca4, can interface with multiple host factors, adding to the growing list of host machinery targeted by these pathogens and highlighting how *Rickettsia* spp. could use their limited genomes to target an even greater number of proteins or pathways. As more rickettsial effectors are identified during infection (73–76), broader effector-interactome studies are warranted in *Rickettsia* spp. to gain a greater appreciation for the scale and reach of additional multifunctional effectors. Complementary approaches investigating the functional consequences of these interactions will also be critical. Such work will deepen our understanding of not only effector biology, but also provide a more complete picture of how pathogens can co-opt their hosts and vectors throughout infection.

## Methods

### Cell Culture

A549 human lung epithelial, HEK293T human embryonic kidney epithelial, and Vero African green monkey kidney epithelial cell lines (University of California, Berkeley Cell Culture Facility, Berkeley, CA) were grown in Dulbecco’s modified Eagle’s medium (DMEM; Gibco 11965118) supplemented with 10% fetal bovine serum (A549, HEK293T) or 5% fetal bovine serum (Vero) at 37°C in a 5% CO2 humidified incubator. *Ixodes scapularis* embryonic-derived ISE6 cells (RRID: CVCL_Z170, ATCC) were maintained in L-15B medium supplemented with 10% heat-inactivated fetal bovine serum (FBS), and 10% tryptose phosphate broth and maintained at 32°C in a 5% CO2 humidified incubator. Cell lines were confirmed to be mycoplasma negative in a MycoAlert PLUS assay (Lonza LT07-710) performed by the Koch Institute High-Throughput Sciences Facility (Cambridge, MA).

### Plasmid Construction

*R. parkeri* Sca4 truncations for IP or pulldown experiments were cloned into pcDNA3 and contain either a 3xFLAG epitope or a GST epitope, followed by a linker (GTAG) and the PCR amplified Sca4 domains, which were codon optimized for human cell expression (32). For *R. typhi* and *R. bellii* Sca4-D1 constructs, JCat (77) was used to codon optimize Sca4 for human expression before Sca4-D1 was synthesized by Twist Biosciences and then cloned into pcDNA3. pRL197 (pcDNA-V5-VCLhead) was made by cloning a PCR amplified human vinculin head domain (1-836aa) into a pcDNA3 with a N-terminal V5 epitope tag (GKPIPNPLLGLDST).

Sca4-D1-D2 constructs were codon optimized for expression in human cells, synthesized by Twist Biosciences, and cloned into pTWIST-CMV with a C-terminal GST epitope tag.

pRAM18dSGA-Sca4pr-Fl-ΔD1Sca4 (pRL329) was created by replacing Sca4 with a PCR amplified product of *R. parkeri* ΔD1Sca4 into pRAM18dSGA-Sca4pr-Fl-Sca4 (32).

### Generation of *R. parkeri* Strains

Parental *R. parkeri* Portsmouth strain (kindly provided by Chris Paddock) and all derived strains were propagated in Vero cells grown in 2% FBS containing DMEM and any appropriate antibiotics for selection at 33°C and isolated through mechanical disruption, centrifugation, and filtering as previously described (32, 78, 79). The *sca4*::Tn, FLAG-Sca4 (*sca4*::Tn + pRAM18dSGA-Sca4pr-Fl-Sca4) and RpGFP (WT + pRAM18dRGA+OmpApr-GFPuv) strains have been previously described (32). The complement, FLAG-Sca4 (*sca4*::Tn + pRAM18dSGA-Sca4pr-Fl-Sca4) and the FLAG-ΔD1-Sca4 (*sca4*::Tn + pRAM18dSGA-Sca4pr-Fl-ΔDSca4) strain were created through electroporation of the *sca4*::Tn mutant (32). Clonal isolation of transformants was performed via plaque purification with 50 µg/mL spectinomycin to select for transformants (32). All bacterial stocks were stored in BHI (Fisher DF0037-17-8) at −80°C.

### Immunoprecipitations

For immunoprecipitations from infected host cell lysates, confluent A549 cell monolayers (approx 3.5 x 10^5^ cells/cm^2^) in 10 cm dishes were infected with WT *R. parkeri* in 800 µl BHI and gently rocked in a humidified chamber at 37°C for 30 min and then incubated at 33°C with 10mL of complete media for 28 h. Cells were washed with 1xPBS and scraped in 1mL IP lysis buffer (50 mM HEPES [pH 7.9], 150 mM NaCl, 1 mM EDTA, 10% glycerol, 1% IGEPAL) supplemented with protease inhibitor cocktail (ApexBio K1008). Lysates were incubated on ice for 25 min before being spun at 11.3K x g for 15 min at 4°C. The supernatant was filtered with a 0.22 µm centrifugation filter (MilliporeSigma UFC30GV00) for 10 min at 4°C at 6,700 x g. Samples were pre-cleared twice for 30 min at 4°C with nProtein A sepharose 4 fast flow (Cytiva 17-5280-01) after which input lysate samples were collected and frozen at −20°C. 1µg of antibody (rabbit anti-Sca4 (32) or mouse anti-CHC (Thermo Fisher Scientific MA1065) was added to remaining lysate and incubated overnight at 4°C. Lysates were then incubated for 1 h at 4°C with 30 µL nProtein A sepharose 4 fast flow and washed 4 times for 5 min each in IP lysis buffer. For mass spectrometry, samples were eluted with 100mM glycine (pH 2.8) and neutralized with 1M Tris (pH 8.5). Otherwise, samples were eluted by boiling for 10 min in 3xSDS loading buffer (150mM Tris-HCl (pH 6.8), 6% SDS, 0.3% Bromophenol Blue, 30% glycerol, 15% BME). Western blot detection used mouse anti-CHC or rabbit anti-Sca4 antibody.

For immunoprecipitations from transfected cells, 5 x 10^5^ HEK-293T cells were plated per well of a 6-well plate. The following day, cells were transfected with 2µg of DNA in a calcium phosphate transfection and the media was changed 22 hours later. Approximately 48 h post-transfection, cells were scraped into 500µL of IP lysis buffer, incubated on ice for 10 min, and then spun at 11.3K x g for 10 min. Supernatants were then pre-cleared twice for 30 min with Protein G sepharose 4 fast flow (Fisher Scientific 45-000-116) at 4°C before either antibody (mouse anti-FLAG, Sigma-Aldrich F1804) or glutathione sepharose was added for overnight incubation. Antibody-treated samples were incubated with sepharose for 1 h at 4°C, then all samples were washed 4 times for 5 min at 4°C before elution by boiling for 10 min in SDS loading buffer. Western blot analysis used the following antibodies: mouse anti-FLAG, rabbit anti-CHC (Abcam ab21679), mouse anti-GST (Santa Cruz sc138), rabbit anti-Sca4 (32), mouse anti-V5 (Invitrogen 46-0707).

### Mass Spectrometry

To identify the Sca4-interacting proteins in the co-IP eluates, S-trap micro spin columns (Protifi) were used according to the manufacturer’s protocol with 10 mM DTT instead of TCEP. Samples were heated for 10 minutes at 95°C then alkylated with 20 mM iodoacetamide and incubated at RT for 30 min in the dark. The tryptic peptides were separated by reverse-phase HPLC (Thermo Ultimate 3000) using a Thermo PepMap RSLC C18 column (2 µm tip, 75 µm x 50 cm PN# ES903) over a 90 min gradient before nano electrospray using an Orbitrap Exploris 480 mass spectrometer (Thermo) in data-dependent mode using standard reverse phase gradients. Initial analysis used Sequest HT in Proteome Discoverer (Thermo) searched against Human (Uniprot) and *Rickettsia parkeri* str. Portsmouth TaxID 1105108 (NCBI) with common contaminants removed from an in-house generated list. Intensities from the top three precursors were calculated in Scaffold Proteomics software. Samples were normalized to the sum of abundance per sample/median sample sum and then proteins were filtered to require a non-zero value for at least two of the three replicates in at least one condition. Remaining zeros were imputed with the half global minimum abundance. Samples were log-transformed and mean fold-changes and adjusted p-values with a two-stage setup (Benjamini, Krieger, and Yekutieli) were calculated.

### Infectious Focus Assays

For infectious focus assays with *CLTC* knockdown, reverse transfections were performed according to the Thermo Fisher RNAiMax reverse transfection protocol. Briefly, 6 pmol of siRNA (siCHC-1: Silencer Select s475, siCHC-2: s477 Thermo Fisher 4390824) were mixed with 100 µl of OptiMEM reduced serum media (Thermo Fisher 31985070) and added to glass coverslips in a 24-well plate along with 1 µl of Lipofectamine RNAiMAX (Thermo Fisher Scientific 13778030). After a 20 min incubation, 1.75 x 10^5^ cells in 0.5mL complete media were added to wells and incubated at 37°C for 48 h. Cells were then infected with an MOI of ~ 0.001, spun for 5 min at 200 x g, and then incubated at 33°C for 1 h. Samples were washed 3 times with PBS and complete media containing 10 µg/mL gentamicin was added. Samples were fixed at 28 hpi with 4% PFA for 15 min.

For infectious focus assays treated with dynasore, A549 cells were grown to confluency (3.5 x 10^5^ cells/cm^2^) on 12mm glass coverslips in 24-well plates. Cells were infected with *R. parkeri* (MOI ~ 0.001), spun at 200 x g for 5 min at RT, and incubated at 33°C for 1 h. Samples were washed 3 times with PBS and complete media containing 10 µg/mL gentamicin was added. At 21 hpi, cells were treated with 40 µM dynasore (Sigma Aldrich D7693) or an equivalent volume of vehicle (ethanol) in 2% serum media for 8 hours. Cells were then fixed with 4% PFA for 15 min.

Fixed samples were stained at RT as follows. Samples were treated with 0.1M glycine for 10 min, washed three times with PBS, and then permeabilized with 0.5% Triton X-100 in PBS for 5 min. Samples were washed twice with PBS and then a blocking buffer (2% bovine serum albumin (BSA) in PBS) was applied for 30 min. Samples were incubated with primary antibody (mouse anti-β-catenin (Cell Signaling 2677); rabbit anti-*Rickettsia* (gift from Ted Hackstadt)) for 1 h and then washed three times for 5 min with PBS. Secondary antibodies (goat anti-rabbit secondary antibody Alexa Fluor-488 (Invitrogen A-11008), goat anti-mouse secondary antibody Alexa Fluor-568 (Invitrogen A-11004) and Hoechst (Invitrogen H3570) were added and incubated for 1 h and followed by three 5 min washes in PBS. Coverslips were mounted in ProLong Gold Antifade (Invitrogen P36934) and imaged with confocal microscopy at 60x magnification (UPlanSApo,1.30 numerical aperture objective) on an Olympus IXplore Spin microscope system with a Yokogawa CSU-W1 spinning-disk unit, ORCA-Flash4.0 sCMOS camera, and Olympus cellSens imaging software. To quantify spread, 20-25 infectious foci per condition were manually analyzed using FIJI software (80).

### Mixed-cell assays

For mixed-cell assays evaluating spread in different *R. parkeri* strains, A549 cells were grown to confluency in 96 well (donor) and 6 well plates (recipient). The donor cell wells were treated with 2 µM CytoTrace Orange CMTMR dye (ATT Bioquest 22014) in serum free media for 20 min at 37°C. Cells were washed with warm PBS and returned to complete media. Cells were infected with an MOI 0.5-1 of indicated strains and incubated for 1 h at 33°C. Both donor and recipient cell samples were washed once with PBS and then treated with 37°C citric saline (135mM KCl, 15mM sodium citrate) for 12 min to gently lift the cells. Single-cell suspensions were recovered in complete media, washed once with complete media to remove citric saline, and then resuspended in complete media containing 10 µg/mL gentamicin (donors cells approximately 2.8 x 10^5^ cells/mL, recipients 2.3 x 10^5^ cells/mL). Cells were then thoroughly mixed at a 1:125 donor:recipient ratio, plated on 12mm glass coverslips in 24 well plates, and incubated at 33°C for a total infection time of 31 h, followed by fixation with 4% PFA in PBS for 45 min at RT.

For mixed-cell assays with *CLTC* knockdown, the manufacturer-suggested reverse transfection protocol described above was scaled for a 6 well plate and used to silence *CLTC* expression in recipient cells. 2 mL of 3.5 x 10^5^ cells/mL were transfected for each recipient cell condition. Donor cells were plated simultaneously in complete media in 96 well plates at 3.5 x 10^4^ cells/well. After cells were incubated at 37°C for 48 hours, the assay proceeded as described above.

For all mixed-cell assays, cells were stained with the same general protocol described in the infectious focus assay with the following antibodies and stains. Rickettsia were stained with mouse anti-*Rickettsia* 14-13 (kindly provided by Ted Hackstadt) and goat anti-mouse Alexa Fluor 488. Phalloidin-647 (Thermo Fisher Scientific A22287) was used to mark actin tails and cell boundaries for ease of quantification. Rabbit anti-CHC (Abcam ab21679) followed by goat anti-rabbit Alex Fluor 405 (Thermo Scientific A-31556) was used to visualize *CLTC* knock-down when appropriate. Cells were imaged as above with >35 foci imaged per condition. Images were processed and manually counted using FIJI (80). Graphs show percent spread as the number of bacteria in recipient cells/total bacteria in the infectious focus.

### Sca4 secretion assay

Sca4 secretion was detected during infection using a selective lysis protocol as previously described (79). Briefly, confluent A549 cells (approx. 3.5 x 10^5^ cells/cm^2^) were infected with the indicated strain of *R. parkeri* for 48 h before being lifted, lysed with IP lysis buffer, and centrifuged to separate the intact bacterial pellet from the supernatant containing host cell material and secreted bacterial effectors. Lysates were analyzed by Western blot using mouse anti-RpoA (BioLegend 663104) and rabbit anti-Sca4.

### Transferrin uptake and recycling assays

For transferrin uptake assays, previously described A549-Sca4 (FCW2-P2AT-Ires-Sca4) or empty vector control (FCW2-P2AT) cells (32) were plated on glass coverslips at 6 x 10^4^ cells/well to allow for well-separated cells. The following day, prewarmed 37°C complete media containing 25 µg/mL human transferrin-CF640 (Biotium 00085) was added to cells. For the 0 min condition, cells were chilled on ice for 15 min before incubating with chilled transferrin-640. After transferrin incubation at 37°C for the indicated time point, all cells were placed on ice and washed twice with cold HBSS + 0.5% BSA. Cells were then acid washed (0.2M acetic acid + 0.5M NaCl) for 2 min on ice before being washed twice with cold HBSS + 0.5% BSA. Cells were then fixed with 4% PFA in PBS for 30 min on ice. For infected transferrin assays, 4 x 10^5^ A549 cells were infected at MOI 1 with RpGFP or *sca4*::Tn for 24 h before cells were lifted with trypsin, diluted 1:10 in complete media, plated on glass coverslips, and incubated at 33°C. At 48 hpi, fluorescent transferrin was added to cells and the assay proceeded as described for the transferrin uptake assay in stable cell lines.

For transferrin recycling assays, cells were plated and treated with transferrin-640 as above and then incubated at 37°C for 1 hour to allow trafficking pathways to become saturated with fluorescent transferrin. Cells were then washed twice with chilled HBSS + 0.5% BSA before 37°C complete media supplemented with 50 µg/mL of unlabeled transferrin was added for the indicated time point. Cells were washed once with HBSS + 0.5% BSA and then fixed with 4% PFA in PBS for 30 min on ice.

For both uptake and recycling assays, fixed cells were treated with 0.1M glycine for 10 min, then washed three times with HBSS. Hoechst and wheat-germ agglutinin (WGA)-CF488 or WGA-CF561 (Biotium) in HBSS were added for 15 min at 37°C. Coverslips were then washed twice in HBSS for 5 min at RT before being mounted in Prolong gold for confocal imaging at 60x. Images were sum projected in FIJI (80) and analysis was completed in CellProfiler 4 (81) by measuring the mean transferrin intensity per cell for >200 cells per condition.

### *R. parkeri* growth kinetics in tick cells

ISE6 cells were seeded onto 24-well plates at a density of 4 x 10^5^ cells/ well and incubated at 32°C for 48 h prior to infection. Purified cell-free rickettsiae of *R. parkeri* wild-type or *R. parkeri sca4*::Tn mutant strains were quantified using a LIVE/DEATH BacLight Bacterial Viability Kit and counted with a Petroff-Hausser bacterial chamber under a fluorescent microscope (69, 82, 83) before being added to ISE6 cells at MOI 1. Samples were centrifuged at 500 x g for 5 minutes at RT and incubated for 1 h at 32°C. At 1 hpi, the unbound bacteria were removed, and these samples served as the initial time point where rickettsial growth was calculated as a change over time. The remaining wells were replaced with L-15B media for a duration of 4 days. Samples containing infected cells, including supernatant, were collected every 24 hours for four days for gDNA extraction and quantification of rickettsial loads by qPCR. A total of 2 independent experiments were performed with three technical replicates per experiment.

All gDNA was extracted via a Qiagen DNeasy blood and tissue kit (Qiagen) following the manufacturer’s instructions. The number of bacterial DNA copies was quantified by qPCR with primers and probes using iTaq Universal Probes Supermix (Bio-Rad) on a LightCycler 480 II (Roche Life Sciences). Each qPCR run included a standard curve of 10-fold serial dilutions of a known concentration of *R. parkeri ompB,* and ISE6 *calreticulin* genes, using previously described primer sets and probes (70, 84). DNA from the samples were amplified along with an environmental DNA extraction control, and water (negative control). The analysis of amplification was conducted with LightCycler 480 software.

### Statistical analysis

Statistical analysis was performed using GraphPad Prism 9 and Prism 10 software. See figure legends for statistical parameters, and significance. All Western blot images are representative of at least two independent experiments.

## Acknowledgments

We would like to thank Eric Spooner at the Whitehead Institute Proteomics Core Facility Richard Schiavoni at the Koch Institute Biopolymers & Proteomics Core Facility for experimental support and advice on mass spectrometry. We are grateful to Tomas Kirchausen for his advice on transferrin uptake assays and to Allen Sanderlin for his assistance with data analysis. This work in the Lamason laboratory is supported by the National Institutes of Health (R01 AI155489 to R.L.L.). B. Sit is supported by a fellowship from the National Institutes of Health (F32AI172121). Work in the Macaluso laboratory is funded by the National Institutes of Health (AI077784 to K.R.M.).

---

**Supplementary Figure 1. Sca4-D1 is not required for the interaction with vinculin head domain**

(**A**) Western blots of V5 and Sca4 following FLAG co-IP in HEK293T cells transiently expressing V5-vinculin head domain (V5-Vinc 1-836aa) and either 3xFLAG-Sca4, 3xFLAG-ΔD1-Sca4, or an empty vector control.

**Supplementary Figure 2. Dynasore treatment reduces transferrin uptake in host cells**

(**A**) Mean fluorescent intensity per field of view of confluent A549 cells after 5 min incubation with fluorescently labeled transferrin. Cells were treated with 40 µM of dynasore or vehicle (EtOH) for 8 hr before application of transferrin to mirror conditions used in Fig 3D, E. Representative experimental replicate with 20-25 analyzed images per condition (two-tailed Mann-Whitney test, ****p < 0.0001)

**Supplementary Figure 3. Sca4 does not alter transferrin uptake or recycling**

(**A**) Fluorescent transferrin uptake assay. Mean fluorescence intensity per cell shown in cells stably expressing either Sca4 or an empty vector control.

(**B**) Fluorescent transferrin uptake assay. Mean fluorescence intensity per cell in uninfected A549 cells versus those infected with RpGFP or *sca4*::Tn. **p < 0.005 for RpGFP vs *sca4*::Tn at 5 min.

(**C**) Fluorescent transferrin recycling assay showing mean fluorescence intensity per cell in cells stably expressing Sca4 or an empty vector control (as described in (A)) For A-C, a representative experiment is provided with each point showing the mean ± SEM for at least 200 individual cells per condition. Kruskal-Wallis test with Dunn’s multiple comparison *post-hoc* test was used to compare samples at each time point.

## References

1. Bhavsar AP, Guttman JA, Finlay BB. 2007. Manipulation of host-cell pathways by bacterial pathogens. Nature 449:827–834.

2. Cornejo E, Schlaermann P, Mukherjee S. 2017. How to rewire the host cell: A home improvement guide for intracellular bacteria. Journal of Cell Biology 216:3931–3948.

3. Kellermann M, Scharte F, Hensel M. 2021. Manipulation of Host Cell Organelles by Intracellular Pathogens. IJMS 22:6484.

4. Finlay BB, Cossart P. 1997. Exploitation of Mammalian Host Cell Functions by Bacterial Pathogens. Science 276:718–725.

5. Ray K, Marteyn B, Sansonetti PJ, Tang CM. 2009. Life on the inside: the intracellular lifestyle of cytosolic bacteria. Nat Rev Microbiol 7:333–340.

6. Raab JE, Hamilton DJ, Harju TB, Huynh TN, Russo BC. 2024. Pushing boundaries: mechanisms enabling bacterial pathogens to spread between cells. Infect Immun e00524–23.

7. Isberg RR, O’Connor TJ, Heidtman M. 2009. The Legionella pneumophila replication vacuole: making a cosy niche inside host cells. Nat Rev Microbiol 7:13–24.

8. Escoll P, Mondino S, Rolando M, Buchrieser C. 2016. Targeting of host organelles by pathogenic bacteria: a sophisticated subversion strategy. Nat Rev Microbiol 14:5–19.

9. Lamason RL, Welch MD. 2017. Actin-based motility and cell-to-cell spread of bacterial pathogens. Current Opinion in Microbiology 35:48–57.

10. Latomanski EA, Newton HJ. 2018. Interaction between autophagic vesicles and the *Coxiella*-containing vacuole requires CLTC (clathrin heavy chain). Autophagy 14:1710– 1725.

11. Poirier V, Av-Gay Y. 2015. Intracellular Growth of Bacterial Pathogens: The Role of Secreted Effector Proteins in the Control of Phagocytosed Microorganisms. Microbiol Spectr 3:3.6.12.

12. Wang G, Xia Y, Cui J, Gu Z, Song Y, Chen YQ, Chen H, Zhang H, Chen W. 2014. The Roles of Moonlighting Proteins in Bacteria. 1. Current Issues in Molecular Biology 16:15–22.

13. Henderson B, Martin A. 2011. Bacterial Virulence in the Moonlight: Multitasking Bacterial Moonlighting Proteins Are Virulence Determinants in Infectious Disease. Infect Immun 79:3476–3491.

14. Matos AL, Curto P, Simões I. 2022. Moonlighting in Rickettsiales: Expanding Virulence Landscape. TropicalMed 7:32.

15. Stein MA, Leung KY, Zwick M, Portillo FG, Finlay BB. 1996. Identification of a Salmonella virulence gene required for formation of filamentous structures containing lysosomal membrane glycoproteins within epithelial cells. Molecular Microbiology 20:151–164.

16. Miao EA, Miller SI. 2000. A conserved amino acid sequence directing intracellular type III secretion by Salmonella typhimurium. Proceedings of the National Academy of Sciences 97:7539–7544.

17. Boucrot E, Beuzón CR, Holden DW, Gorvel J-P, Méresse S. 2003. Salmonella typhimurium SifA Effector Protein Requires Its Membrane-anchoring C-terminal Hexapeptide for Its Biological Function. Journal of Biological Chemistry 278:14196–14202.

18. McEwan DG, Richter B, Claudi B, Wigge C, Wild P, Farhan H, McGourty K, Coxon FP, Franz-Wachtel M, Perdu B, Akutsu M, Habermann A, Kirchof A, Helfrich MH, Odgren PR, Van Hul W, Frangakis AS, Rajalingam K, Macek B, Holden DW, Bumann D, Dikic I. 2015. PLEKHM1 Regulates Salmonella-Containing Vacuole Biogenesis and Infection. Cell Host & Microbe 17:58–71.

19. Ohlson MB, Huang Z, Alto NM, Blanc M-P, Dixon JE, Chai J, Miller SI. 2008. Structure and Function of Salmonella SifA Indicate that Its Interactions with SKIP, SseJ, and RhoA Family GTPases Induce Endosomal Tubulation. Cell Host & Microbe 4:434–446.

20. Merhej V, Royer-Carenzi M, Pontarotti P, Raoult D. 2009. Massive comparative genomic analysis reveals convergent evolution of specialized bacteria. Biol Direct 4:13.

21. Kelkar YD, Ochman H. 2013. Genome Reduction Promotes Increase in Protein Functional Complexity in Bacteria. Genetics 193:303–307.

22. Niu H, Kozjak-Pavlovic V, Rudel T, Rikihisa Y. 2010. Anaplasma phagocytophilum Ats-1 Is Imported into Host Cell Mitochondria and Interferes with Apoptosis Induction. PLoS Pathog 6:e1000774.

23. Niu H, Xiong Q, Yamamoto A, Hayashi-Nishino M, Rikihisa Y. 2012. Autophagosomes induced by a bacterial Beclin 1 binding protein facilitate obligatory intracellular infection. Proc Natl Acad Sci USA 109:20800–20807.

24. Walker DH, Ismail N. 2008. Emerging and re-emerging rickettsioses: endothelial cell infection and early disease events. Nat Rev Microbiol 6:375–386.

25. Parola P, Paddock CD, Socolovschi C, Labruna MB, Mediannikov O, Kernif T, Abdad MY, Stenos J, Bitam I, Fournier P-E, Raoult D. 2013. Update on Tick-Borne Rickettsioses around the World: a Geographic Approach. Clin Microbiol Rev 26:657–702.

26. McGinn J, Lamason RL. 2021. The enigmatic biology of rickettsiae: recent advances, open questions and outlook. Pathogens and Disease 79:ftab019.

27. Sit B, Lamason RL. 2024. Pathogenic *Rickettsia* spp. as emerging models for bacterial biology. J Bacteriol 206:e00404–23.

28. Gillespie JJ, Kaur SJ, Rahman MS, Rennoll-Bankert K, Sears KT, Beier-Sexton M, Azad AF. 2014. Secretome of obligate intracellular *Rickettsia*. FEMS Microbiol Rev n/a-n/a.

29. Salje J. 2021. Cells within cells: Rickettsiales and the obligate intracellular bacterial lifestyle. Nat Rev Microbiol 19:375–390.

30. Schuenke KW, Walker DH. 1994. Cloning, sequencing, and expression of the gene coding for an antigenic 120-kilodalton protein of Rickettsia conorii. Infect Immun 62:904–909.

31. Park H, Lee JH, Gouin E, Cossart P, Izard T. 2011. The Rickettsia Surface Cell Antigen 4 Applies Mimicry to Bind to and Activate Vinculin. Journal of Biological Chemistry 286:35096–35103.

32. Lamason RL, Bastounis E, Kafai NM, Serrano R, Del Álamo JC, Theriot JA, Welch MD. 2016. Rickettsia Sca4 Reduces Vinculin-Mediated Intercellular Tension to Promote Spread. Cell 167:670–683.e10.

33. Sears KT, Ceraul SM, Gillespie JJ, Allen ED, Popov VL, Ammerman NC, Rahman MS, Azad AF. 2012. Surface Proteome Analysis and Characterization of Surface Cell Antigen (Sca) or Autotransporter Family of Rickettsia typhi. PLoS Pathogens 8:e1002856.

34. Blanc G, Ngwamidiba M, Ogata H, Fournier P-E, Claverie J-M, Raoult D. 2005. Molecular Evolution of Rickettsia Surface Antigens: Evidence of Positive Selection. Molecular Biology and Evolution 22:2073–2083.

35. Silverman DJ, Wisseman CL. 1979. In vitro studies of rickettsia-host cell interactions: ultrastructural changes induced by Rickettsia rickettsii infection of chicken embryo fibroblasts. Infect Immun 26:714–727.

36. Brodsky FM. 2012. Diversity of Clathrin Function: New Tricks for an Old Protein. Annu Rev Cell Dev Biol 28:309–336.

37. Johnson RP, Craig SW. 1995. F-actin binding site masked by the intramolecular association of vinculin head and tail domains. Nature 373:261–264.

38. Weiss EE, Kroemker M, Rüdiger A-H, Jockusch BM, Rüdiger M. 1998. Vinculin Is Part of the Cadherin–Catenin Junctional Complex: Complex Formation between ␣-Catenin and Vinculin. The Journal of Cell Biology 141.

39. Bakolitsa C, Cohen DM, Bankston LA, Bobkov AA, Cadwell GW, Jennings L, Critchley DR, Craig SW, Liddington RC. 2004. Structural basis for vinculin activation at sites of cell adhesion. Nature 430:583–586.

40. Poon IKH, Patel KK, Davis DS, Parish CR, Hulett MD. 2011. Histidine-rich glycoprotein: the Swiss Army knife of mammalian plasma. Blood 117:2093–2101.

41. Kumar M, Michael S, Alvarado-Valverde J, Mészáros B, Sámano-Sánchez H, Zeke A, Dobson L, Lazar T, Örd M, Nagpal A, Farahi N, Käser M, Kraleti R, Davey NE, Pancsa R, Chemes LB, Gibson TJ. 2022. The Eukaryotic Linear Motif resource: 2022 release. Nucleic Acids Research 50:D497–D508.

42. Dell’Angelica EC. 2001. Clathrin-binding proteins: Got a motif? Join the network! Trends in Cell Biology 11:315–318.

43. Bryson K, Cozzetto D, Jones DT. 2007. Computer-assisted protein domain boundary prediction using the DomPred server. Curr Protein Pept Sci 8:181–188.

44. Lee JH, Vonrhein C, Bricogne G, Izard T. 2013. Crystal structure of the *N*-terminal domains of the surface cell antigen 4 of *Rickettsia*. Protein Science 22:1425–1431.

45. Chan YGY, Cardwell MM, Hermanas TM, Uchiyama T, Martinez JJ. 2009. Rickettsial outer-membrane protein B (rOmpB) mediates bacterial invasion through Ku70 in an actin, c-Cbl, clathrin and caveolin 2-dependent manner. Cellular Microbiology 11:629–644.

46. Macia E, Ehrlich M, Massol R, Boucrot E, Brunner C, Kirchhausen T. 2006. Dynasore, a Cell-Permeable Inhibitor of Dynamin. Developmental Cell 10:839–850.

47. Fukumatsu M, Ogawa M, Arakawa S, Suzuki M, Nakayama K, Shimizu S, Kim M, Mimuro H, Sasakawa C. 2012. Shigella Targets Epithelial Tricellular Junctions and Uses a Noncanonical Clathrin-Dependent Endocytic Pathway to Spread Between Cells. Cell Host & Microbe 11:325–336.

48. Mayle KM, Le AM, Kamei DT. 2012. The intracellular trafficking pathway of transferrin. Biochimica et Biophysica Acta (BBA) - General Subjects 1820:264–281.

49. Dautry-Varsat A. 1986. Receptor-mediated endocytosis: The intracellular journey of transferrin and its receptor. Biochimie 68:375–381.

50. Li J, Peters PJ, Bai M, Dai J, Bos E, Kirchhausen T, Kandror KV, Hsu VW. 2007. An ACAP1-containing clathrin coat complex for endocytic recycling. The Journal of Cell Biology 178:453–464.

51. Hsu VW, Bai M, Li J. 2012. Getting active: protein sorting in endocytic recycling. Nat Rev Mol Cell Biol 13:323–328.

52. Azad A. 1998. Rickettsial Pathogens and Their Arthropod Vectors. Emerg Infect Dis 4:179–186.

53. Perlman SJ, Hunter MS, Zchori-Fein E. 2006. The emerging diversity of *Rickettsia*. Proc R Soc B 273:2097–2106.

54. Humphries AC, Way M. 2013. The non-canonical roles of clathrin and actin in pathogen internalization, egress and spread. Nat Rev Microbiol 11:551–560.

55. Latomanski EA, Newton HJ. 2019. Taming the Triskelion: Bacterial Manipulation of Clathrin. Microbiol Mol Biol Rev 83:e00058–18.

56. Cureton DK, Massol RH, Saffarian S, Kirchhausen TL, Whelan SPJ. 2009. Vesicular Stomatitis Virus Enters Cells through Vesicles Incompletely Coated with Clathrin That Depend upon Actin for Internalization. PLoS Pathog 5:e1000394.

57. Bhattacharyya S, Warfield KL, Ruthel G, Bavari S, Aman MJ, Hope TJ. 2010. Ebola virus uses clathrin-mediated endocytosis as an entry pathway. Virology 401:18–28.

58. Veiga E, Cossart P. 2005. Listeria hijacks the clathrin-dependent endocytic machinery to invade mammalian cells. Nat Cell Biol 7:894–900.

59. Moreno-Ruiz E, Galán-Díez M, Zhu W, Fernández-Ruiz E, d’Enfert C, Filler SG, Cossart P, Veiga E. 2009. Candida albicans internalization by host cells is mediated by a clathrin-dependent mechanism. Cell Microbiol 11:1179–1189.

60. Nagajyothi F, Weiss LM, Silver DL, Desruisseaux MS, Scherer PE, Herz J, Tanowitz HB. 2011. Trypanosoma cruzi Utilizes the Host Low Density Lipoprotein Receptor in Invasion. PLoS Negl Trop Dis 5:e953.

61. Larson CL, Beare PA, Howe D, Heinzen RA. 2013. *Coxiella burnetii* effector protein subverts clathrin-mediated vesicular trafficking for pathogen vacuole biogenesis. Proc Natl Acad Sci USA 110.

62. Latomanski EA, Newton P, Khoo CA, Newton HJ. 2016. The Effector Cig57 Hijacks FCHO-Mediated Vesicular Trafficking to Facilitate Intracellular Replication of Coxiella burnetii. PLoS Pathog 12:e1006101.

63. Yuan Y, Wang X, Jin J, Tang Z, Xian W, Zhang X, Fu J, He K, Liu X. 2023. The Salmonella Typhimurium Effector SpvB Subverts Host Membrane Trafficking by Targeting Clathrin and AP-1. Molecular & Cellular Proteomics 22:100674.

64. Royle SJ, Lagnado L. 2006. Trimerisation is important for the function of clathrin at the mitotic spindle. Journal of Cell Science 119:4071–4078.

65. Royle SJ, Bright NA, Lagnado L. 2005. Clathrin is required for the function of the mitotic spindle. Nature 434:1152–1157.

66. Deborde S, Perret E, Gravotta D, Deora A, Salvarezza S, Schreiner R, Rodriguez-Boulan E. 2008. Clathrin is a key regulator of basolateral polarity. Nature 452:719–723.

67. McCormick PJ, Martina JA, Bonifacino JS. 2005. Involvement of clathrin and AP-2 in the trafficking of MHC class II molecules to antigen-processing compartments. Proc Natl Acad Sci USA 102:7910–7915.

68. Kim ML, Sorg I, Arrieumerlou C. 2011. Endocytosis-Independent Function of Clathrin Heavy Chain in the Control of Basal NF-κB Activation. PLoS ONE 6:e17158.

69. Tongluan N, Engström P, Jirakanwisal K, Langohr IM, Welch MD, Macaluso KR. 2024. Critical roles of *Rickettsia parkeri* outer membrane protein B (OmpB) in the tick host. Infect Immun e00515–23.

70. Harris EK, Jirakanwisal K, Verhoeve VI, Fongsaran C, Suwanbongkot C, Welch MD, Macaluso KR. 2018. Role of Sca2 and RickA in the Dissemination of Rickettsia parkeri in Amblyomma maculatum. Infect Immun 86:e00123–18.

71. McMahon HT, Boucrot E. 2011. Molecular mechanism and physiological functions of clathrin-mediated endocytosis. Nat Rev Mol Cell Biol 12:517–533.

72. Kuang C, Wang F, Zhou Y, Cao J, Zhang H, Gong H, Zhou R, Zhou J. 2020. Molecular characterization of clathrin heavy chain (Chc) in Rhipicephalus haemaphysaloides and its effect on vitellogenin (Vg) expression via the clathrin-mediated endocytic pathway. Exp Appl Acarol 80:71–89.

73. Sanderlin AG, Margolis HK, Meyer AF, Lamason RL. 2023. Cell-selective proteomics reveal novel effectors secreted by an obligate intracellular bacterial pathogen 10.1101/2023.11.17.567466.

74. Voss OH, Gillespie JJ, Lehman SS, Rennoll SA, Beier-Sexton M, Rahman MS, Azad AF. 2020. Risk1, a Phosphatidylinositol 3-Kinase Effector, Promotes Rickettsia typhi Intracellular Survival. mBio 11:e00820–20.

75. Lehman SS, Noriea NF, Aistleitner K, Clark TR, Dooley CA, Nair V, Kaur SJ, Rahman MS, Gillespie JJ, Azad AF, Hackstadt T. 2018. The Rickettsial Ankyrin Repeat Protein 2 Is a Type IV Secreted Effector That Associates with the Endoplasmic Reticulum. mBio 9:e00975–18.

76. Rennoll-Bankert KE, Rahman MS, Gillespie JJ, Guillotte ML, Kaur SJ, Lehman SS, Beier-Sexton M, Azad AF. 2015. Which Way In? The RalF Arf-GEF Orchestrates Rickettsia Host Cell Invasion. PLoS Pathog 11:e1005115.

77. Grote A, Hiller K, Scheer M, Munch R, Nortemann B, Hempel DC, Jahn D. 2005. JCat: a novel tool to adapt codon usage of a target gene to its potential expression host. Nucleic Acids Research 33:W526–W531.

78. Paddock CD, Sumner JW, Comer JA, Zaki SR, Goldsmith CS, Goddard J, McLellan SLF, Tamminga CL, Ohl CA. 2004. *Rickettsia parkeri:* A Newly Recognized Cause of Spotted Fever Rickettsiosis in the United States. CLIN INFECT DIS 38:805–811.

79. Sanderlin AG, Hanna RE, Lamason RL. 2022. The Ankyrin Repeat Protein RARP-1 Is a Periplasmic Factor That Supports *Rickettsia parkeri* Growth and Host Cell Invasion. J Bacteriol 204:e00182–22.

80. Schindelin J, Arganda-Carreras I, Frise E, Kaynig V, Longair M, Pietzsch T, Preibisch S, Rueden C, Saalfeld S, Schmid B, Tinevez J-Y, White DJ, Hartenstein V, Eliceiri K, Tomancak P, Cardona A. 2012. Fiji: an open-source platform for biological-image analysis. Nat Methods 9:676–682.

81. Stirling DR, Swain-Bowden MJ, Lucas AM, Carpenter AE, Cimini BA, Goodman A. 2021. CellProfiler 4: improvements in speed, utility and usability. BMC Bioinformatics 22:433.

82. Laukaitis HJ, Cooper TT, Suwanbongkot C, Verhoeve VI, Kurtti TJ, Munderloh UG, Macaluso KR. 2022. Transposon mutagenesis of Rickettsia felis sca1 confers a distinct phenotype during flea infection. PLoS Pathog 18:e1011045.

83. Sunyakumthorn P, Bourchookarn A, Pornwiroon W, David C, Barker SA, Macaluso KR. 2008. Characterization and Growth of Polymorphic Rickettsia felis in a Tick Cell Line. Applied and Environmental Microbiology 74:3151–3158.

84. Banajee KH, Embers ME, Langohr IM, Doyle LA, Hasenkampf NR, Macaluso KR. 2015. Correction: Amblyomma maculatum Feeding Augments Rickettsia parkeri Infection in a Rhesus Macaque Model: A Pilot Study. PLOS ONE 10:e0137598.

